# Non-linear changes in modelled terrestrial ecosystems subjected to perturbations

**DOI:** 10.1101/439059

**Authors:** Tim Newbold, Derek P. Tittensor, Michael B. J. Harfoot, Jörn P. W. Scharlemann, Drew W. Purves

## Abstract

When perturbed ecosystems undergo rapid and non-linear changes, this can result in ‘regime shifts’ to an entirely different ecological state. The need to understand the extent, nature, magnitude and reversibility of these changes is urgent given the profound effects that humans are having on the natural world. It remains very challenging to empirically document non-linear changes and regime shifts within complex, real ecological communities, or even to demonstrate such shifts in simplified experimental systems. General ecosystem models, which simulate the dynamics of entire ecological communities based on a mechanistic representation of ecological processes, provide an alternative and novel way to project ecosystem changes across all scales and trophic levels and to forecast impact thresholds beyond which dramatic or irreversible changes may occur. We model non-linear changes in four terrestrial biomes subjected to human removal of plant biomass, such as occurs through agricultural land-use change. We find that irreversible and non-linear responses are predicted to be common where removal of vegetation exceeds 80% (a level that occurs across nearly 10% of the terrestrial surface), especially for organisms at higher trophic levels and in less productive ecosystems such as drylands. Very large, irreversible changes to the entire ecosystem structure are expected at levels of vegetation removal akin to those in the most intensively used real-world ecosystems. Our results suggest that the projected 21^st^ century rapid increases in agricultural land conversion to feed an expanding human population, may lead to widespread trophic cascades and in some cases irreversible changes to the structure of ecological communities.

---

Human activities are impacting ecosystems across the globe, leading to impaired ecosystem function (Hooper et al. 2012), altered trophic structure (Branch et al. 2010), trophic cascades (Frank et al. 2005; Pace et al. 1999) and – with sufficient levels of impact – ecosystem collapse (MacDougall et al. 2013). Such changes can have serious consequences for the human societies that rely on those ecosystems (Millennium Ecosystem Assessment 2005). Because ecosystems are very complex, involving many interacting entities, it is likely that responses to change do not occur linearly, and ecosystems under increasing pressure may eventually undergo rapid regime shifts to fundamentally different states (Scheffer et al. 2001; Estes et al. 2011). For example, coral reefs subject to human disturbance can shift rapidly to an algal-dominated state (Scheffer et al. 2001; Mumby, Hastings, and Edwards 2007), and loss of top predators can lead to marked shifts in vegetation structure (Estes et al. 2011). However, in many cases these changes are not well understood, or predictable, particularly in terms of the threshold of impact beyond which a transition will occur. Ecosystems undergoing rapid non-linear changes or regime shifts can exhibit hysteresis, where the trajectory followed by the ecosystem subsequent to alleviation of a pressure differs from that during escalation of the pressure (Scheffer et al. 2001). Changes may also be irreversible, with removal of the pressure not leading to a full recovery of the system, and possibly leading to the establishment of an alternative state (Scheffer et al. 2001).

The search for empirical evidence of ecosystem responses to perturbations has tended to use simplified experimental systems, or to focus on particular ecosystems and on incomplete subsets of the species in those ecosystems, often top predators, large herbivores or vegetation (Scheffer et al. 2001; Estes et al. 2011). However, there are more complete empirical studies of a few well-studied systems (Frank et al. 2005; Carpenter et al. 2011). An alternative way to explore perturbation responses is to simulate the effects of perturbations using ecological models of the structure of complex ecological communities. Widely used statistical models (Newbold et al. 2015; Warren et al. 2013) cannot generally capture or predict non-linear and dynamic responses of whole ecosystem structure to perturbations (Evans et al. 2013; but see e.g. Wearn, Reuman, and Ewers 2012), in part because they do not explicitly model the complex interplay of processes underlying the functioning of ecosystems. Alternatively, models of specific subsets of the food web in particular ecosystems have been used to explore how human perturbations might impact particular species or groups of species, and ultimately cause the ecosystem subset to collapse (Dakos and Bascompte 2014; Dunne and Williams 2009). However, these models do not come close to capturing the whole range of organisms within the ecosystem.

In contrast, mechanistic ecosystem models (Purves et al. 2013; Fulton et al. 2011; Villy Christensen and Walters 2004; Harfoot et al. 2014) simulate the underlying biological interactions among individual organisms, and other processes structuring ecosystems. Therefore, these models can begin to capture complex and dynamic interactions, and so are, in principle, much better suited to predicting dynamical changes and novel configurations in whole ecosystems. By using a general ecosystem model that represents all plants and non-microbial heterotrophic organisms within ecosystems (as ‘functional groups’, within which organisms are assumed to play a similar role in the ecosystem), the age- and size-structuring of ecosystems, processes such as metabolism and growth, predator-prey interactions, and spatial interactions between different locations, we ask whether collapses are likely to occur in more complex ecological systems. This approach also allows us to apply the same general ecosystem model to four very different ecosystems.

We use the Madingley Model (Harfoot et al. 2014), a general ecosystem model, to simulate human impacts on the fundamental structure of a complex terrestrial ecosystem, including all autotrophs and all heterotrophs larger than 10 μg. In sample ecosystems within four terrestrial biomes, spanning the major global productivity and seasonality gradients (Table 1), we ask how ecosystem-level properties – including both total biomass and functional properties – respond to the imposition of human perturbations, and in a second set of simulations we explore how ecosystems respond to the subsequent removal of human perturbation. Specifically we ask: 1) whether responses to perturbations of broad-scale ecosystem properties (biomasses and abundance) are non-linear; 2) whether high-level functional properties of ecosystem also respond to perturbation; 3) whether ecosystems recover to pre-perturbation states, or whether impacts lead to alternative states after recovery.

**Table 1.** Locations and characteristics of the ecosystems simulated.

| Country | Longitude | Latitude | Productivity | Seasonality | Panel letter in figures |
| --- | --- | --- | --- | --- | --- |
| Uganda | 33 | 2 | Very High | Low | a |
| France | 4 | 48 | Moderately high | High | b |
| China | 91 | 38 | Moderately low | High | c |
| Libya | 13 | 30 | Very low | Low | d |

Land use is one of the primary human perturbations to terrestrial biodiversity globally (Newbold et al. 2015), and removal of plant biomass is one of the main manifestations of human use of the land, both as a result of converting natural vegetation to human use, and subsequently harvesting plant biomass from agricultural areas (Haberl et al. 2007). We used removal of plant biomass, expressed as a fraction of net primary production (NPP), as a proxy for the impacts of land use on ecosystems. Analysis focuses on uniform perturbations to ecosystems across a 3 × 3 grid at 1° spatial resolution, and does not consider landscape-wide effects of habitat fragmentation that have been considered elsewhere (Bartlett et al. 2016).

As expected, as levels of NPP removal increased, the biomass in all trophic levels and the abundance of herbivores, omnivores and carnivores were all reduced (Figures 1 & 2). The maximum level of NPP removal currently experienced by real-world ecosystems is 94% (Haberl et al. 2007). At this level of human pressure, changes to ecosystem structure in the modelled system were profound, with losses of biomass and abundance greater than 60% in all modelled ecosystems, and approaching 100% losses in many cases (Figures 1 & 2). The functional properties of ecosystems also responded strongly to perturbation (Figure 3). The ranges of both body masses and trophic levels present in the ecosystem declined substantially (Figure 3), supporting suggestions that human impacts are simplifying the structure of ecosystems (Estes et al. 2011). The trophic level reduction was driven entirely by the loss of organisms at high trophic levels, i.e. top predators, because plants and herbivores were never completely lost from ecosystems (Appendix 1, Figure S2); for body mass range both the largest and smallest organisms were often lost, although the pattern varied across simulated ecosystems (Appendix 1, Figure S3). As suggested previously in empirical studies, for subsets of organisms within ecosystems (Estes et al. 2011; Pauly et al. 1998), mean trophic level also declined (Figure 3).

**Figure 1.**
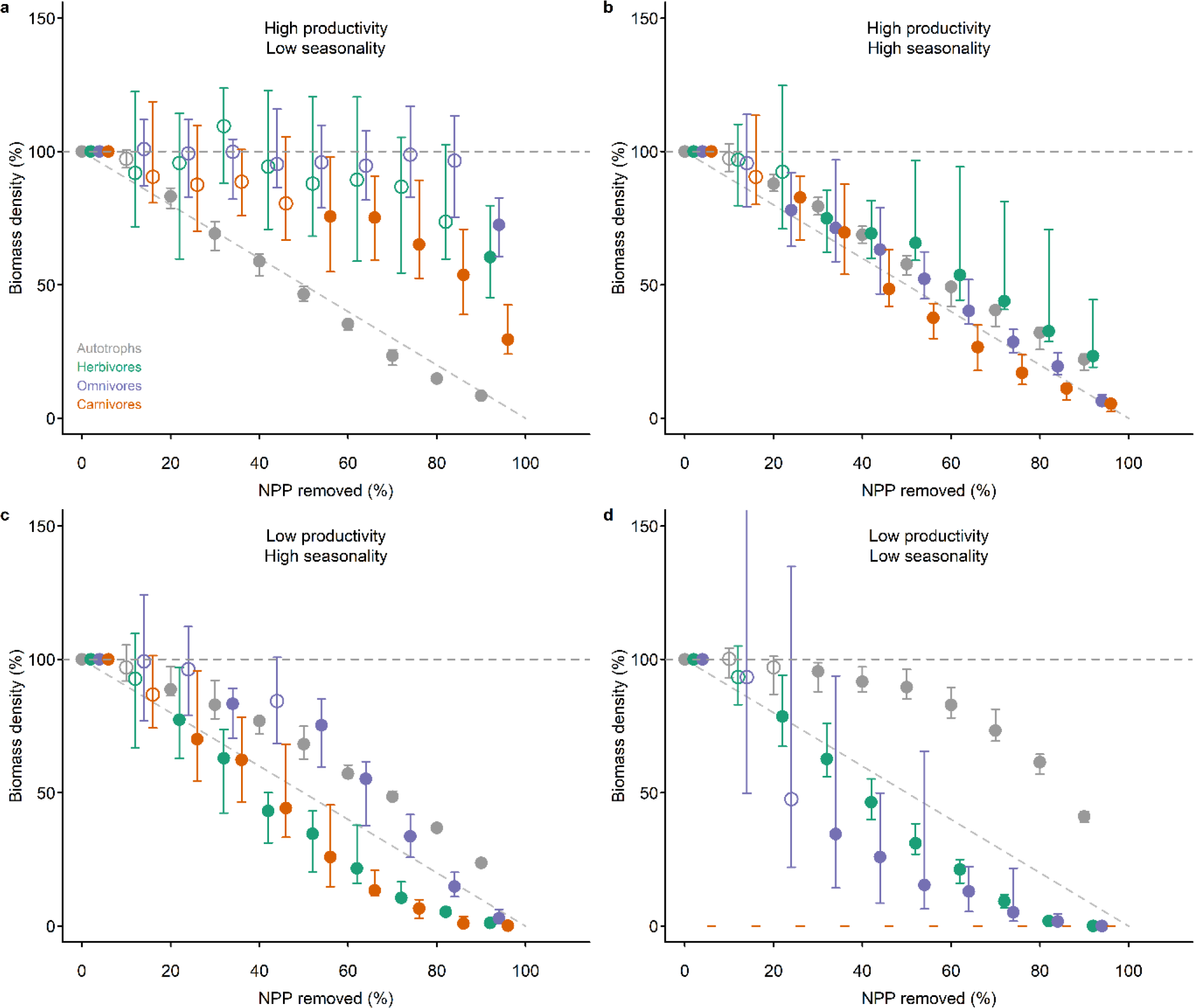
Impact, relative to an undisturbed state, of the removal of net primary production (NPP) on the biomass of organisms in different trophic levels ‒ autotrophs (grey), herbivores (green), omnivores (purple) and carnivores (red) ‒ in four ecosystems: (a) high productivity, aseasonal (Uganda); (b) relatively high productivity, highly seasonal (France); (c) relatively low productivity, seasonal (Gobi Desert, China); and (d) low productivity, aseasonal (Libyan Desert). Ecosystems were subjected to levels of NPP removal that increased by 1% per year for between 0 and 90 years, resulting in maximum levels of NPP removal ranging between 0 and 90%. Values shown here are the ecosystem properties after 100 years since the onset of impact. Error bars show ±95% confidence intervals; open points show effects with confidence intervals overlapping 100%. The grey, horizontal dashed line indicates projected values for unimpacted ecosystems. The diagonal dashed line indicates the impact expected if that impact occurred in direct proportion to the amount of plant productivity removed (i.e. y = −x).

**Figure 2.**
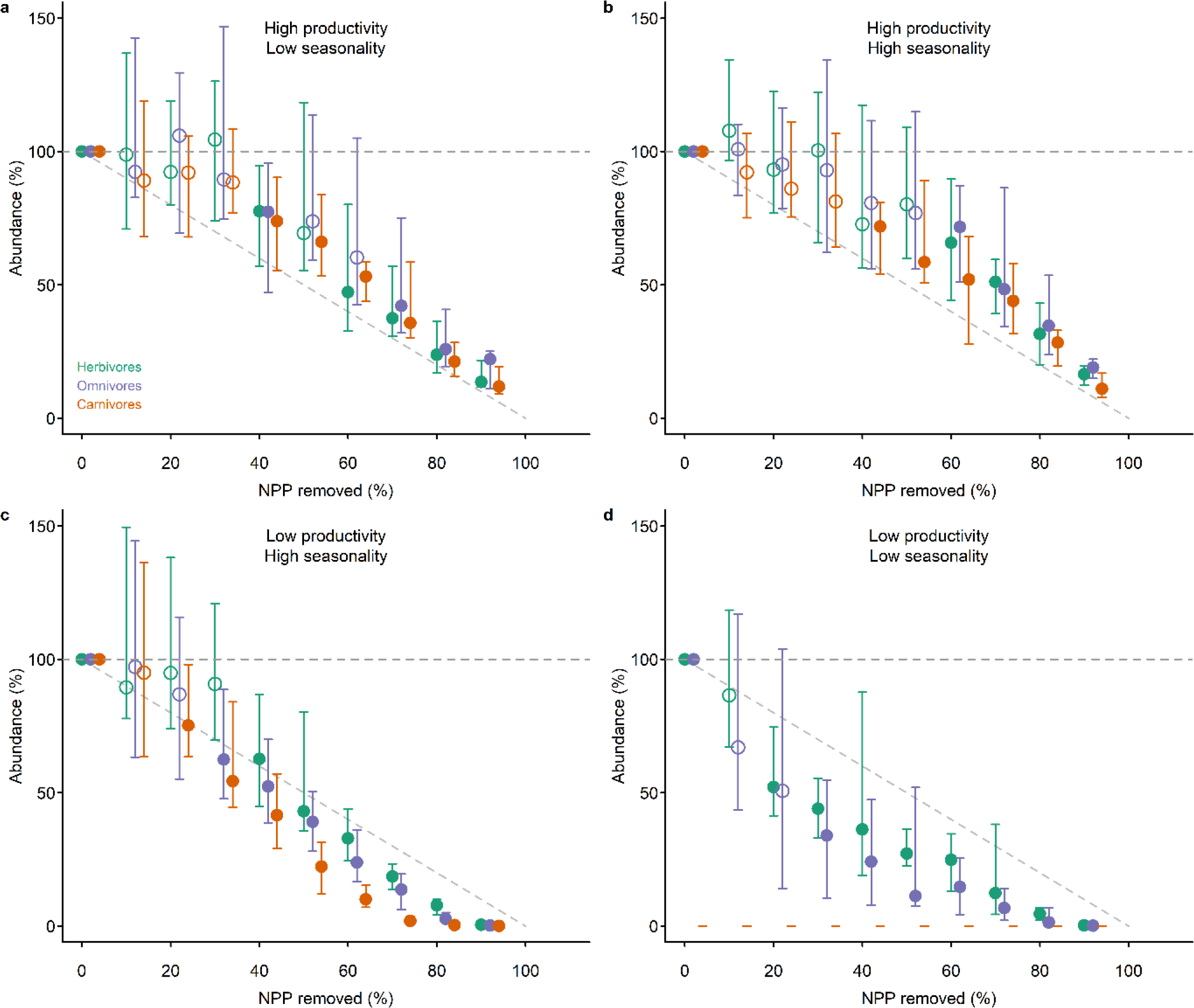
Impact, relative to an undisturbed state, of removal of net primary production (NPP) on the abundance of organisms in different trophic levels ‒ herbivores (green), omnivores (purple) and carnivores (red) ‒ in four ecosystems: (**a**) high productivity, aseasonal (Uganda); (**b**) relatively high productivity, highly seasonal (France); (**c**) relatively low productivity, seasonal (Gobi Desert, China); and (**d**) low productivity, aseasonal (Libyan Desert). Simulations and plotting conventions are as in Figure 1.

**Figure 3.**
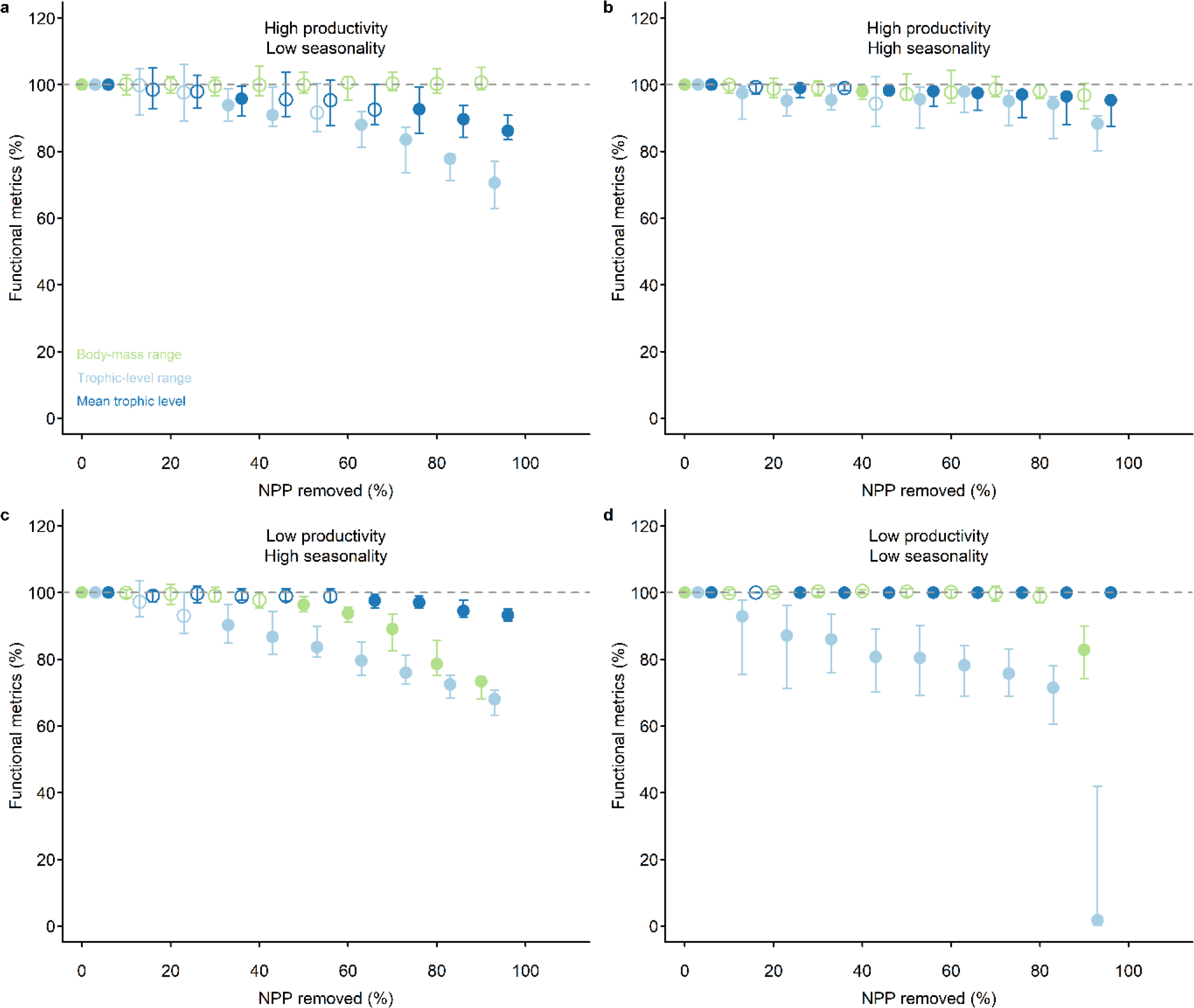
Impact, relative to an undisturbed state, of removal of net primary production (NPP) on the functional structure ‒ mean trophic level (dark blue), trophic range (light blue) and body-mass range (light green) ‒ of four ecosystems: (**a**) high productivity, aseasonal (Uganda); (**b**) relatively high productivity, highly seasonal (France); (**c**) relatively low productivity, seasonal (Gobi Desert, China); and (**d**) low productivity, aseasonal (Libyan Desert). Simulations and plotting conventions are as in Figure 1.

In most cases responses to human pressure were non-linear (shown by departures from the dashed line at y = −x in Figures 1 & 2). The non-linearity was more pronounced: 1) for biomass than for abundance; 2) for organisms in the higher trophic levels; and 3) for the two locations with low seasonality, whether of high (Figure 1a) or low overall productivity (Figure 1d). The disproportionate non-linearity in the responses of high trophic levels is likely because of the dynamic nature of predator-prey interactions and because the resources for predators are scarcer and more patchily distributed in nature, making predators more sensitive to bottom-up resource limitation. In the high-productivity, low-seasonality system, non-linear responses to perturbation occurred because higher trophic levels persisted until a high proportion of plant biomass was removed. This system was characterized by a very low natural ratio (0.15%) of heterotrophs to autotrophs (Appendix 1, Figure S4), meaning that top-down forces are likely more important in structuring the system than bottom-up resource limitation, at least under conditions of low plant biomass removal. In the low-productivity, low-seasonality (desert) system, non-linear responses occurred because heterotrophs were lost very rapidly, even at low levels of perturbation. This system had a very high ratio (8.9%) of heterotrophs to autotrophs, and very low plant biomass, placing a high degree of resource limitation, and thus sensitivity to resource reduction, on higher trophic levels. The impact threshold at which the most rapid changes occurred (in terms of both level of human pressure and the intactness of the ecosystem prior to the rapid changes) varied depending on location and on the property measured (compare panels a-d in Figures 1-3). For example, in the highly productive and aseasonal tropical-forest system, rapid changes in ecosystem structure occurred only for levels of vegetation removal greater than 50%, whereas for the lower-productivity systems, rapid changes occurred even at very low levels of vegetation removal (Figures 1 & 2).

The non-linear behaviour of ecosystems predicted here is supported by empirical evidence suggesting that gradual increases in levels of impact can lead to abrupt shifts between alternative stable states (Scheffer et al. 2001). However, our results go further in suggesting the existence of threshold levels of impact (varying here from less than 50% to greater than 90% removal of vegetation biomass, depending on the ecosystem simulated) beyond which ecosystems undergo almost complete collapse to a state of greatly reduced complexity and diversity, and minimal biomass. Importantly, levels of removal of plant biomass currently experienced in the most intensively used ecosystems (94% of primary productivity removed by humans; Haberl et al. 2007) are on the edge of such thresholds.

Under the highest observed levels of human pressure, changes to ecosystem structure were substantial and often irreversible (Figure 4). Gradually increasing vegetation removal to 95% and then gradually reducing it back to zero, led to temperate forest, temperate arid and desert ecosystems that were substantially altered compared to before perturbation. Irreversibility was more likely for higher trophic levels and in less productive ecosystems (Figure 4). This is likely because landscape-wide biomasses were more likely to reach very low levels for high trophic levels and in less productive ecosystems (Figure 1), with a corresponding reduction in functional diversity (Figure 3), making rescue effects from immigration much less likely. Irreversibility was less pronounced, but still present, for lower levels of removal of plant biomass – up to 80%, or up to 90% (Appendix 1, Figures S5 & S6) – levels of plant-biomass removal already estimated to occur across 9.4% and 0.03% of the terrestrial area, respectively (Haberl et al. 2007).

**Figure 4.**
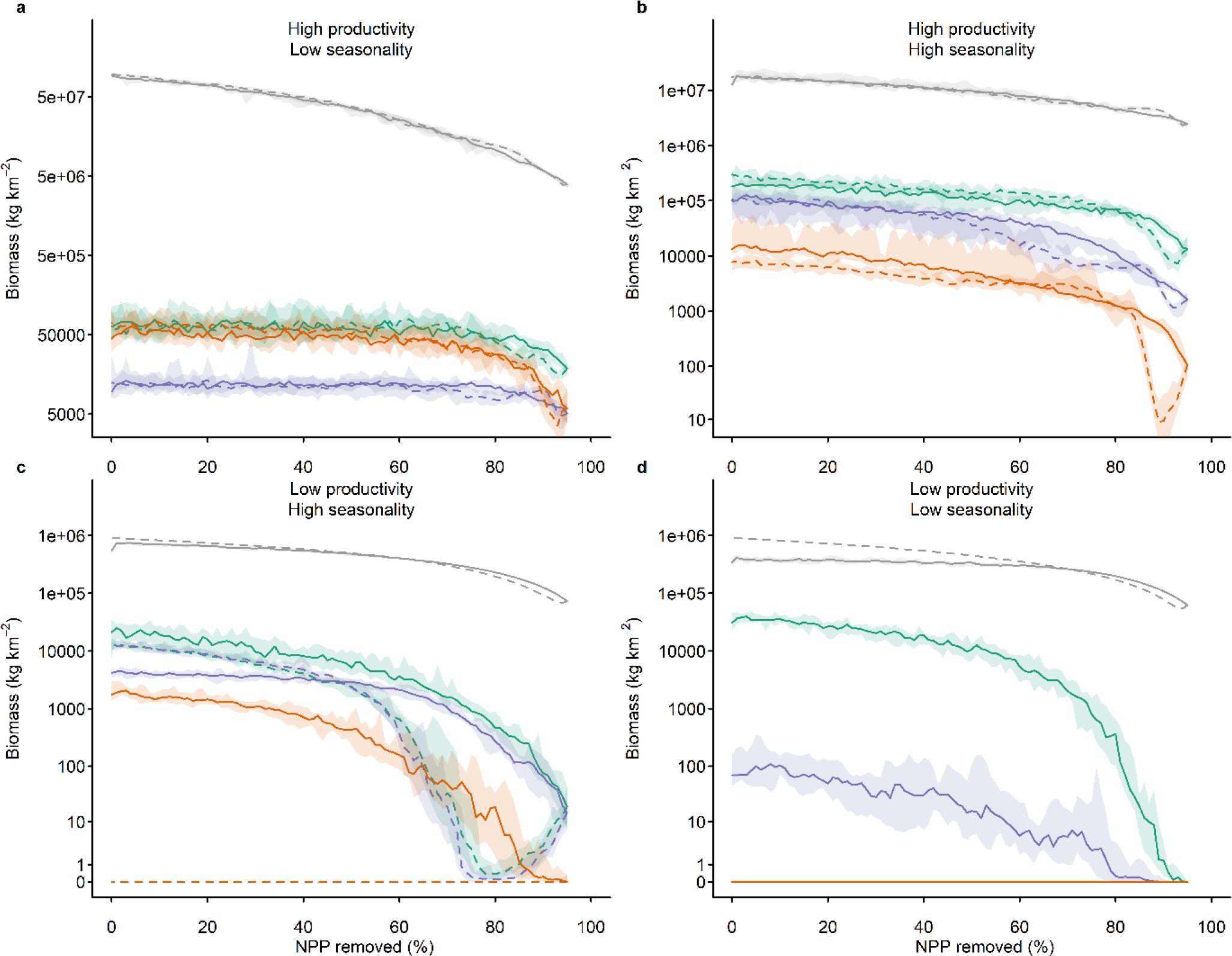
Irreversibility of change in total ecosystem biomass during a gradual increase in human impact (solid lines), for 95 years by 1% of total primary production per annum to 95% removal; followed by a gradual release (dotted lines), by 1% per annum to zero impact. Lines show biomasses in different trophic levels ‒ autotrophs (grey), herbivores (green), omnivores (purple) and carnivores (red) in four ecosystems: (**a**) high productivity, aseasonal (Uganda); (**b**) relatively high productivity, highly seasonal (France); (**c**) relatively low productivity, seasonal (Gobi Desert, China); and (**d**) low productivity, aseasonal (Libyan Desert). Shading shows 95% confidence intervals.

The irreversibility of human impacts is relevant to ongoing discussions about ecosystem restoration (Seddon et al. 2014), suggesting that without external interventions restored ecosystems may be unlikely to approach the structure or function of their natural state. The context-dependency of the irreversibility suggests that extrapolating effects observed for specific ecosystems and ecosystem components to other situations may give misleading results, and that further ecosystem-specific analysis is necessary to understand the full suite of idiosyncrasies and contingencies associated with each biome that ultimately inform its susceptibility to irreversible regime shifts.

Ecosystem complexity, unprecedented simultaneous impacts upon them and limited empirical evidence for non-linear responses and regime shifts in real ecosystems has led to problems in identifying, at a global scale, important thresholds of ecosystem change beyond which increasingly rapid changes are expected. A recent attempt to find a ‘planetary boundary’ for biodiversity, beyond which the whole earth system might depart from the relatively stable state characteristic of the Holocene, suggested a threshold value of somewhere between 30 and 90% of the total number of individuals remaining in ecosystems compared with the pre-perturbation state (Steffen et al. 2015). The very wide uncertainty range makes the proposed boundary of limited use for policy application. Statistical modelling has suggested that the precautionary upper limit to this planetary boundary has been surpassed across much of the Earth’s land surface (Newbold et al. 2016). This result, combined with the observation that most ecosystems have not ceased functioning, suggests either that lagged effects are likely to manifest in the future, or that the upper limit is overly conservative.

Our results suggest that NPP removal pushes ecosystems towards collapse, while fragmentation reduces the potential for rescue (as shown elsewhere: Bartlett et al. 2016). As more of the Earth’s land surface becomes highly impacted by humans, it is possible that local or regional shifts will combine, and potentially cascade, to lead to global regime shifts in the Earth’s biosphere (Rockström et al. 2009). Our results point toward large spatial variation in the location of regime shifts, which will be important to predict potential global thresholds, and suggests that mapping such variation in thresholds should become an urgent concern. We also show similar variation in the degree of ecosystem transformation, often irreversible, for a given level of NPP removal, which has profound implications for establishing ‘safe operating spaces’, levels of perturbation below which the risk of destabilising ecosystems and the services they provide is likely to remain low. This variability suggests that defining a ‘planetary boundary’ for biodiversity in terms of the intactness of ecosystems (Steffen et al. 2015) will be subject to a high degree of contingency and uncertainty, or at the very least will need to be defined in terms of regional specifics and ecosystem-to-ecosystem and biome-to-biome variation.

There are a number of caveats to the analyses. First, for computational tractability we focused on four ecosystems. These ecosystems were selected to represent a set of the major terrestrial climatic gradients. Further work would have to be conducted to determine the applicability and transferability of the reported patterns to other systems. Second, comprehensive data on ecosystem structure with which to evaluate the predictions of general ecosystem models are generally lacking. However, tests against available data suggest that the Madingley Model captures many properties of ecosystems reasonably well (Harfoot et al. 2014). Third, our ecosystems were simulated in isolation (though with a grid of 3 × 3 1° cells), and thus without the possibility for rescue by immigration from adjacent less-disturbed areas (although note that rescue from adjacent disturbed areas was possible). In many parts of the world there are few such areas from which rescue effects could occur, as they are undergoing a combination of increased removal of NPP at local scales (Krausmann et al. 2013), and habitat fragmentation across the wider landscape, which can have important and deleterious effects on biodiversity (Fahrig 2003; Krauss et al. 2010). Our simulations also excluded the possibility for long-distance dispersal to rescue populations following human disturbance, which can be important (Brooker et al. 2007). Fourth, obligate carnivores were predicted to be absent from the simulated desert ecosystem even without human impacts.

This erroneous prediction probably arises because the Madingley Model does not represent behavioural processes that allow predators to exploit scarce food resources, through long-distance migration or disproportionately large home range sizes, aggregation around patches of denser food supply, and dormancy. Fifth, model experiments to disentangle possible mechanisms for the patterns reported here will be an interesting avenue to explore in future, but beyond the scope of the current study.

In conclusion, most attempts to predict the future of ecological communities have focused on species-centred measures of diversity (Jetz, Wilcove, and Dobson 2007; Visconti et al. 2016), and it has been suggested that loss of species could impair ecosystem function (Hooper et al. 2012). Here we provide a first-principles demonstration of how human impacts are expected to change the structure of whole ecosystems and their functioning, given the complexities that structure those ecosystems. We demonstrate a profound expected impact of the removal of net primary production (NPP) – one of the most important ways that humans influence the biosphere (Haberl et al. 2007) – on the fundamental structure of ecosystems. We demonstrate that ecosystems are expected to show non-linear responses to human perturbations, undergoing a rapid collapse at high but realistic levels of impact. Under more moderate levels of impact, ecosystems are not expected to collapse, but nonetheless undergo major shifts in functional structure. After release from impact, ecosystems cannot be expected to recover to their pre-perturbation state, at least in the absence of external rescue effects.

Defining a safe operating space for biodiversity will therefore be difficult, as modelled ecosystems appear to be irreversibly transformed substantially before levels of perturbation that would cause collapse, and levels of impact leading to substantial ecosystem changes varied strongly among ecosystems. Fundamentally, our results suggest that the predicted increases in NPP appropriation by humans in the coming century (Krausmann et al. 2013) will cause profound and potentially irreversible ecosystem changes, with unknown but almost certainly negative consequences for humanity. The complexity of real ecosystems is not necessarily sufficient to buffer them against collapse.

## Methods

### The Madingley Model

We simulated ecosystems in the Madingley Model. The Madingley Model is a mechanistic General Ecosystem Model that simulates the dynamics of all photoautotrophs, and all heterotrophs with body masses above 10μg that feed on living organisms (Harfoot et al. 2014). Organisms are characterized by a combination of functional traits rather than by species identity. Categorical traits are used to group organisms into a set of functional groups – for example, leaf strategy and mobility for autotrophic organisms; trophic level (herbivores, omnivores and carnivores), reproductive strategy (semelparity vs. iteroparity), thermoregulatory mode (endothermy vs. ecothermy) and mobility for heterotrophic organisms. Continuous traits – total biomass of autotrophs; and current body mass, juvenile body mass, adult body mass, and optimal prey size of heterotrophs – also determine the outcome of ecological processes. The full source code for the Madingley Model with all necessary input data can be downloaded freely at https://github.com/Madingley. Alternatively, a pre-compiled version – ready to run the simulations and with all input data – can be downloaded at http://dx.doi.org/10.6084/m9.figshare.6531194.

Autotrophic organisms are represented as stocks of plant biomass, and the temporal change of these stocks is modelled through the processes of primary productivity, growth and mortality (including from herbivory). Heterotrophic animals are represented in the model as cohorts, which are collections of individual organisms occurring in the same modelled grid cell with identical categorical and very similar continuous functional traits. We could not represent individual organisms for computational reasons (Purves et al. 2013) but this approach enabled the model to predict emergent properties of individuals (Harfoot et al. 2014). Performance of animals and interactions among them are captured in the ecological processes of eating, metabolism, reproduction, mortality and dispersal. The ecological processes of autotrophs and heterotrophs occur within, and are influenced by, the environment. For the terrestrial realm, which was the focus of this study, the environment is represented by air temperature, precipitation, soil water availability, number of frost days, and seasonality of primary productivity. The model can be run at varying spatial and temporal resolutions, but has been tested principally with 1° and 2° grid cells, and with time steps of one month (Harfoot et al. 2014). Within each grid cell, the environment is assumed to be spatially homogeneous. This is an important limitation of the model in its current form.

### Perturbation Simulations

At each of four locations (Table 1), we subjected a 3 × 3 grid of 1° × 1° grid cells to different levels of vegetation removal. Removal of vegetation was simulated by allowing a proportion of the primary production in a given cell to be appropriated for human uses: this was defined as a proportion of primary production rather than a proportion of total plant biomass, for comparability with global datasets (Haberl et al. 2007). Primary production was calculated as a function of climate (Harfoot et al. 2014). Appropriated biomass was assumed to become entirely unavailable to organisms in the modelled ecosystems.

Vegetation removal was applied to the model following two protocols. Ten replicates of both protocols at each level of human impact were applied to each of ten simulated ecosystems, which had first been run for 1,000 years with no human impacts to allow dynamic steady-state ecosystems to emerge. All simulations were run with a time step of one month. At the end of the no-impact initialization simulations the model states for each of the ten replicate ecosystems was saved and used as input to the human-impact simulations. Contemporary monthly climate data were used throughout all simulations. Temperature, precipitation, and diurnal temperature range data were derived from Microsoft Research’s FetchClimate tool (Grechka et al. 2016). Available soil water capacity estimates were taken from a global dataset (ISRIC-WISE 2012).

The first protocol (Appendix 1, Figure S1a), which was used for most of the results presented, involved gradually increasing the level of pressure (1% increase per year) until different specified levels of impact were reached, ranging from 0% removal to 90% removal of primary production. Once the specified level was reached, the ecosystem was subjected to this level of impact until a total of 100 years had elapsed (from the point at which the escalation of impact began), after which the impact was removed and the ecosystem was given the opportunity to recover for a further 100 years.

The second protocol (Appendix 1, Figure S1b) was used only to test whether ecosystems recovered to the pre-perturbation state after gradual removal of human pressure (i.e. whether responses to human perturbations were reversible). This protocol involved gradually increasing the level of pressure (1% increase per year) until 95% of primary production was removed (just above the maximum level currently experienced by any terrestrial ecosystem; Haberl et al. 2007). We also repeated these simulations with maximum rates of removal of 80% and 90% of primary production, to test the sensitivity of the results. Once these maximum levels were reached, the impact was gradually decreased (by 1% per year) back to zero impact. These simulations were used to test whether ecosystems showed irreversible responses to impact.

We simulated four locations with different natural environmental conditions (Table 1): 1) high productivity, aseasonal (tropical forest in Uganda); 2) relatively high productivity, seasonal (temperate forest in Central France); 3) relatively low productivity, seasonal (Gobi Desert, China); and 4) low productivity, aseasonal (Libyan Desert). The same level of impact was applied to all nine grid cells in the simulated grid.

### Output Metrics

At each monthly time-step, we calculated the following measures of ecosystem structure and functional composition: biomass of autotrophs, herbivores, omnivores and carnivores; abundance of herbivores, omnivores and carnivores; the range of body masses and trophic levels, and the mean trophic level of organisms present.

We calculated the trophic index of each cohort (collections of organisms occurring in the same grid cell and with similar traits) as a continuous value based on its trophic interactions. Using an adaptation of the equation used by Christensen and Pauly (Christensen and Pauly 1992) we calculated the trophic index at time step *t* for a cohort *i* (*T_i_*) based on the mean trophic index of all prey items weighted by the dietary fraction each represents for *i*:

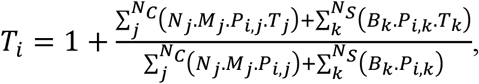

where *j* is a prey cohort, from among all cohorts *N*_*C*_ in the grid cell; *N*_*j*_ is the abundance, *M*_*j*_ the total mass (including reproductive potential mass) and *T*_*j*_ the trophic index of prey cohort *j*; *T*_*i,j*_ is the fraction of cohort *j* eaten by predator cohort *i*; *k* is an autotroph stock eaten, from among all stocks *N*_*S*_ in the grid cell; *B*_*k*_ is the biomass and *T*_*k*_ the trophic index of the stock (*T*_*k*_ = 1); *P*_*i,k*_ is the fraction of stock *k* eaten by cohort *i*. For the calculation of the fraction of prey cohorts and plant stocks eaten, and parameters for the relevant equations, see (Harfoot et al. 2014).

The mean trophic level was calculated as the arithmetic mean trophic index across all cohorts weighted by the total biomass in each cohort:

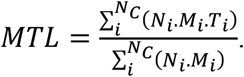

Trophic-level range at time *t* was calculated as the difference between the maximum and minimum trophic indices across all cohorts *i* in a grid cell, standardized by dividing by the difference between a hypothetical maximum (*T*_*max*_ = 40.0) and the minimum possible (*T*_*min*_ = 1.0) trophic indices:

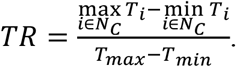

Body-mass range at time *t* was similarly calculated as the difference between the maximum and minimum body masses across all cohorts *i* in a grid cell, standardized by dividing by the difference between the maximum (*M*_*max*_) and minimum (*M*_*min*_) potential body masses allowed in the model (0.00001 g and 150,000,000 g, respectively):

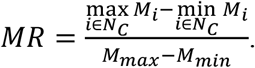

There is a README file within both of the model downloads (links given above) that explains the practicalities of running the simulations with the Madingley Model. This information is also given in Appendix 2.

## Acknowledgements

This work was supported by the UN Environment World Conservation Monitoring Centre, Microsoft Research, a Leverhulme Trust Research Project grant (TN), and funding from the Kanne Rassmussen Foundation, Denmark (DPT, MBJH). We thank Matthew J. Smith for valuable discussions about this work, and Neil D. Burgess for comments on the manuscript.

## Author Contributions

T.N., D.P.T., M.B.J.H. and D.W.P. designed the study. T.N. and M.B.J.H carried out the analyses. T.N., D.P.T., M.B.J.H., J.P.W.S. and D.W.P. wrote the manuscript. T.N., D.P.T. and M.B.J.H. contributed equally to the study.

